# Linking the genetic structure of neuroanatomical phenotypes with psychiatric disorders

**DOI:** 10.1101/2023.11.01.564329

**Authors:** Antoine Auvergne, Nicolas Traut, Léo Henches, Lucie Troubat, Arthur Frouin, Christophe Boetto, Sayeh Kazem, Hanna Julienne, Roberto Toro, Hugues Aschard

**Affiliations:** Génétique Statistique, Institut Pasteur, Université de Paris-Cité, Paris, France; U5 Neuroanatomie Appliquée et Théorique, Institut Pasteur, Paris, France

## Abstract

There is increasing evidence of shared genetic factors between psychiatric disorders and brain magnetic resonance imaging (MRI) phenotypes. However, deciphering the joint genetic architecture of these outcomes has proven challenging, and new approaches are needed to infer potential genetic structure underlying those phenotypes. Here, we demonstrate how multivariate analyses can help reveal links between MRI phenotypes and psychiatric disorders missed by univariate approaches. We first conducted univariate and multivariate genome-wide association studies (GWAS) for eight MRI-derived brain volume phenotypes in 20K UK Biobank participants. We performed various enrichment analyses to assess whether and how univariate and multitrait approaches can distinguish disorder-associated and non-disorder-associated variants from six psychiatric disorders: bipolarity, attention-deficit/hyperactivity disorder (ADHD), autism, schizophrenia, obsessive-compulsive disorder, and major depressive disorder. Univariate MRI GWAS displayed only negligible genetic correlation with psychiatric disorders at all the levels we investigated. Multitrait GWAS identified multiple new associations and showed significant enrichment for variants related to both ADHD and schizophrenia. We further clustered top associated variants based on their MRI multitrait association using an optimized *k*-medoids approach and detected two clusters displaying not only enrichment for association with ADHD and schizophrenia, but also consistent direction of effects. Functional annotation analyses pointed to multiple potential mechanisms, suggesting in particular a role of neurotrophin pathways on both MRI and schizophrenia. Altogether our results show that multitrait association signature can be used to infer genetically-driven latent MRI variables associated with psychiatric disorders, opening paths for future biomarker development.

## Introduction

Psychiatric disorders present strong heterogeneity in their clinical presentation and identifying biomarkers remains a critical research area to facilitate diagnosis and implement more focused treatment options^1^. The study of neuroimaging phenotypes and their association with mental disorders has arisen as a major direction to develop robust and reliable biomarkers and pathological feature^2^. MRI phenotypes have the potential to capture endophenotypes related to the molecular and cellular mechanisms involved in the sources of the disorders, and thus improve our understanding of the underlying pathophysiology^3^. Thousands of studies have been published in the field, with approximately a third based on the analysis of structural MRI data (e.g. nuclei volumes and shapes, grey and white matter ratio, etc)^4^. These works have contributed to the identification of links between brain regions and psychiatric disorders, such as the increased ventricular size in schizophrenia^5^. However, other associations have not been consistently identified and remain controversial, with for example, both higher^6^ and lower^7^ putamen volume having been associated with higher risk for schizophrenia.

Neuroimaging genomics is a recent field that offers new opportunities to address these questions through the integration of genomic and imaging data^8^. Regional volumetric phenotypes have a strong genetic component with estimated heritability (*h*^2^) commonly above 50%^9–11^. Many psychiatric disorders, such as schizophrenia (*h*^2^=80%)^5^, bipolar disorder (*h*^2^=80%)^12,13^, attention-deficit/hyperactivity disorder (*h*^2^=74%)^14^, autism spectrum disorder (*h*^2^=64-91%)^15^, obsessive-compulsive disorder (*h*^2^=27-65%)^16^ or major depressive disorder (*h*^2^=32-37%)^17,18^ also display high heritability. Over the past years there has been increasing evidences of genetic correlation between those disorders and MRI phenotypes, shedding new light on potential mechanisms between MRI phenotypes and mental disorders^10,19–25^. However, deciphering the joint genetic architecture of high-dimensional neuroimaging outcomes and how they impact mental disorders has proven challenging, and new approaches are needed for inferring potential shared genetic structure underlying these disorders.

Here, we investigated the potential of multivariate genetic approaches to infer links between MRI phenotypes and mental disorders. As a proof of concept, we considered eight MRI phenotypes already investigated in previous studies^26^ and six disorders: attention-deficit/hyperactivity disorder (ADHD), autism spectrum disorder (ASD), bipolar disorder (BIP), major depressive disorder (MDD), obsessive-compulsive disorder (OCD) and schizophrenia (SCZ). We first conducted univariate and multivariate genome-wide association screenings (GWAS) of the eight MRI phenotypes and examined genetic correlation with each of the six disorders. We then investigated the utility of univariate and multivariate genetic results to improve detection of genetic variants associated with the disorders. Finally, we applied a clustering approach on the genetic results matrix and searched for latent MRI variables displaying consistent genetic signal with mental disorders. This analysis identified candidate latent variables for schizophrenia and ADHD, demonstrating the potential of multivariate MRI genetic analysis to infer potential biomarkers for mental disorders.

## Results

### Univariate and multitrait GWAS of MRI phenotypes

We first assessed the potential gain of using multivariate versus univariate association tests for the analysis of eight MRI phenotypes: volume of the nucleus accumbens, amygdala, caudate nucleus, pallidum, putamen, thalamus, hippocampus and intracranial volume (ICV). We conducted univariate GWAS on 11,993,198 variants using Plink2^27^ (**Fig. S1**) in 20,744 unrelated participants from the UK Biobank^28^. The multivariate analysis was conducted using an omnibus test applied to the univariate summary statistics, as implemented in the JASS software^29^. The univariate GWAS detected a total of 41 genome-wide significant phenotype-locus associations (*P* < 6.25 x 10^-9^ after accounting for the eight GWAS conducted), involving 34 loci, and with six loci displaying significant association for at least two phenotypes (**Table S1**). We compared these results against previous GWAS of the same phenotypes conducted in independent samples as part of the ENIGMA consortium^30^. Five out of the eight associations reported in the ENIGMA study were also genome-wide significant in our analysis, the three remaining being within the threshold of replicability (*P* < 6.2 x 10^-3^, **Table S2**). Out of the 41 associations detected only in the present analysis, we observed a strong enrichment for association in ENIGMA, with 40% (16 out of 41) of the closest top variants being nominally significant (*P* < 0.05) (**Table S1**).

Multitrait analysis detected a total of 49 loci associated at genome-wide significance level (*P* < 5 x10^-8^), corresponding to a 44% increase in power as compared to univariate analysis. This included 22 out of the 34 loci identified with the univariate analysis, and 25 new associations not identified by the univariate screening (**Table S3, Fig. S2**). Minimum univariate *p*-value at these new signals ranged from 0.003 to 1.8 x 10^-8^ highlighting the ability of the multitrait test to detect association with limited single-trait signal. The univariate association pattern varied substantially across these new signals, with some variants displaying modest association with multiple traits, and other displaying strong association with only one or two traits. Notably, we observed enrichment for association with caudate and putamen volumes with almost two third (15 out of 25) of univariate *p*-values being nominally significant. Out of the 25 new associated variants, half of them could be mapped with genes using data available on the NCBI website^31^. GeneCards^32^ search showed that seven of them are associated with brain or psychiatric disorders (**Table S3**).

### Shared genetic factors between neuroanatomical traits and psychiatric disorders

We estimated the genetic correlation between the MRI phenotypes and six disorders, attention-deficit/hyperactivity disorder (ADHD), autism spectrum disorder (ASD), bipolar disorder (BIP), major depressive disorder (MDD), obsessive-compulsive disorder (OCD) and schizophrenia (SCZ) (**Table S4**), using both genome-wide variants based on the LDscore regression^33,34^ and top associated variants from the multitrait test. Genome-wide correlations between disorders were all significant, ranging from 0.27 to 0.71 (schizophrenia and bipolar disorder) (**Fig. 1a**). Similar genetic correlation estimates between these two disorders were obtained by Ruderfer et al^35^, Bulik-Sullivan et al^34^ and others^23^. We observed similarly high correlation across MRI phenotypes, ranging from 0.22 to 0.67 ^26^. Conversely, there was low or no correlations between the MRI phenotypes and the disorders, with few significative correlations ranging from -0.17 (between ADHD and ICV) to 0.24 (between obsessive-compulsive disorder and accumbens volume). Correlation between disorders based on top variants were qualitatively similar (**Fig. 1b**) or slightly lower, with an exception for autism spectrum disorder and attention-deficit hyperactivity disorder, being more significant than genome-wide correlation (*ρ*= 0.77, *P < 1 x 10*^-16^). Correlations between phenotypes at top associated variants were substantially larger than those measured at the genome-wide level, and highly significant, ranging from 0.71 to 0.94 (*P < 1 x 10*^-16^). Few significant correlations were detected between disorders and phenotypes, with the maximum correlation observed between hippocampal volume and ADHD (*ρ* = -0.09, *P* = *1*.*7 x 10*^-4^).

**Figure 1.**
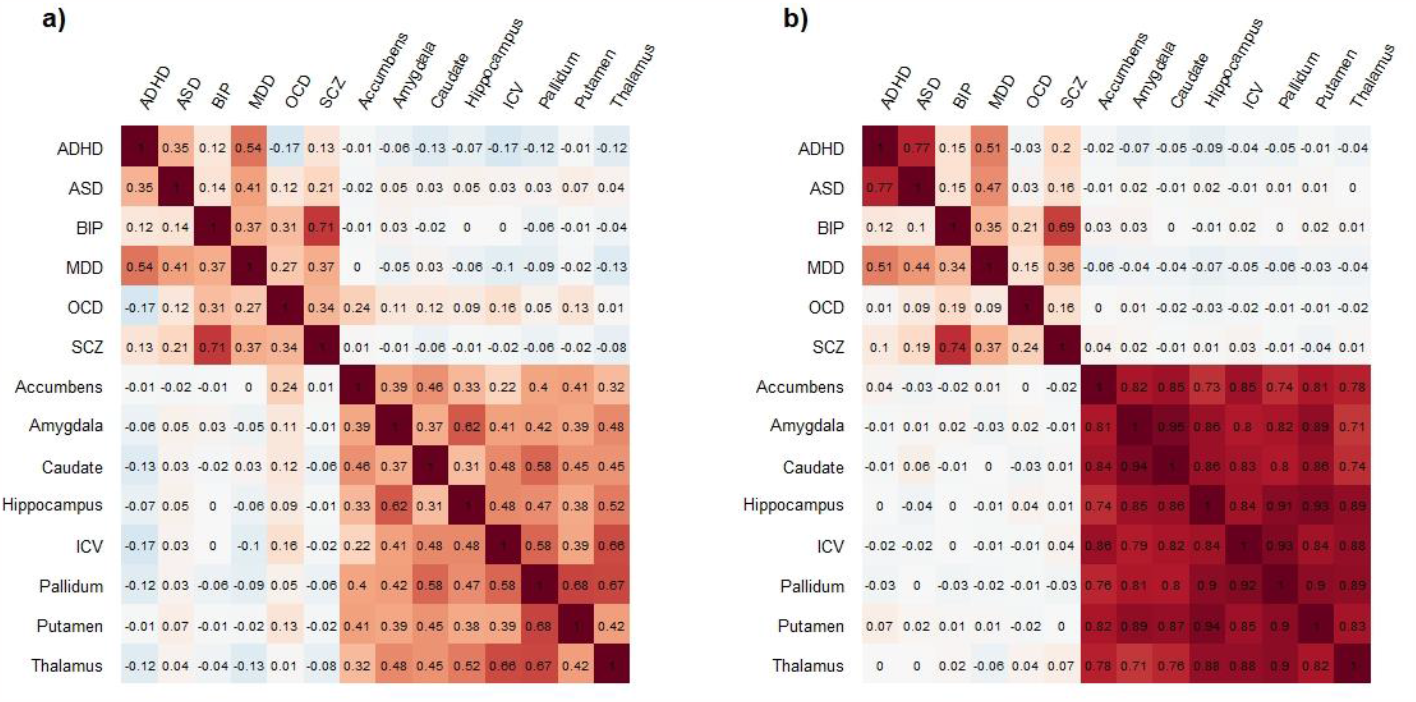
Genome-wide and independent variants correlation. Correlation between eight neuroanatomical phenotypes (accumbens, amygdala, caudate, hippocampus, intracranial volume (ICV), pallidum, putamen, and thalamus) and six disorders: attention-deficit/hyperactivity disorder (ADHD), autism spectrum disorder (ASD), bipolarity (BIP), major depressive disorder (MDD), obsessive-compulsive disorder (OCD) and schizophrenia (SCZ). Correlation was estimated genome-wide using the LDscore regression (a) and using independent genome-wide significant variants (*P* < 5 x10^-8^, b). For independent variants correlation, the correlation matrix is asymmetrical, and variants have been selected based on their association with the phenotype displayed on the line name.

Correlation analyses are based on signed statistics. However, some variants might be associated with both mental disorder and MRI without displaying concordant effect across the genome. To address this question, we assessed whether genetic associations *p*-value with MRI phenotype can inform associations with mental disorders. For each univariate MRI GWAS and for the multitrait analysis, we conducted a stepwise filtering of associated loci based on the strength of significance and tested the enrichment of the top variant per locus with each of the six psychiatric disorders. Note that the size of the disorder’s GWASs implies an overall enrichment of signal for any random set of variants, and simple enrichment analysis method was therefore not applicable. Instead, we used an empirical approach, estimating the probability of enrichment from the selected variants as compared to a null distribution derived through permutation (see **methods**). Except for hippocampus and autism spectrum disorder, which displayed an enrichment just above the significance threshold after correction for multiple testing (*P*_threshold_ = 5.4 x 10^-6^), variants selected through univariate neuroanatomical GWAS results did not show significant association with disorders (**Fig. 2**). Conversely, the variants selected through the multivariate analysis displayed highly significant associations with ADHD and schizophrenia (*P* < 10^-7^, the smallest *p*-value achievable in our permutation-based analysis). For ADHD, enrichment became significant when including variants with *P* < 1.5 x 10^-4^ but was not significant anymore when including only the top associated JASS variants (*P* < 1.5 x 10^-8^), likely because of a limited number of variants in this category (N < 40). For schizophrenia, enrichment became significant when including variants with *P* < 2.9 x 10^-3^ and remained significant under the smallest *p*-value we can detect.

**Figure 2.**
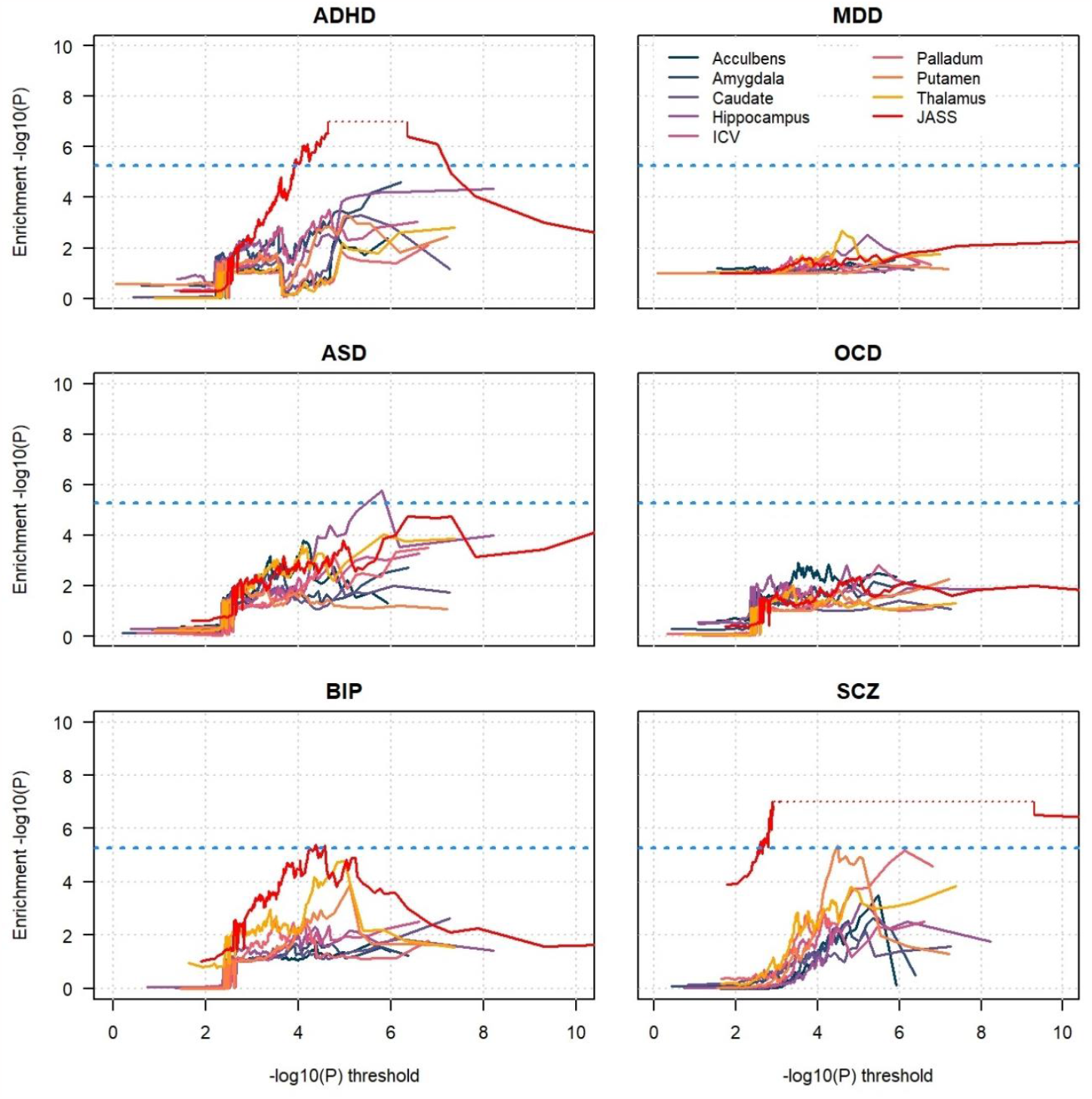
Enrichment for genetic association with disorders conditional on brain phenotype association. We estimated to what extent genetic signal observed with each of the eight neuroanatomical phenotypes were enriched for genetic association with six disorders: attention-deficit/hyperactivity disorder (ADHD), autism spectrum disorder (ASD), bipolarity (BIP), major depressive disorder (MDD), obsessive-compulsive disorder (OCD) and schizophrenia (SCZ). Each panel presents the -log10(*p*-value) for enrichment for a given disorder for 170 subsets of independent genetic variants selected based on their association with the neuroanatomical phenotype. We applied the same approach using the *p*-value from the multitrait association test (JASS, red curve). *p*-value for enrichment were derived using a permutation approach, which limited the minimum *P* that could be reach. Dash red lines represent this maximum, indicating that the empirical *p*-value is smaller than 10^-7^. The dash blue line represent a stringent Bonferroni corrected significant threshold.

As a complementary analysis, we investigated whether this significant enrichment for signal could be translated into improved polygenic risk models using the multitrait results, and schizophrenia as a case study. We derived two polygenic risk scores (PRS), one using a standard approach applied to the schizophrenia GWAS only, and a weighted PRS as implemented in SBayesRC^36^ using the JASS association signal as weight (see **methods**). We applied both PRSs to an independent set of 585 schizophrenia cases and 9,396 controls from the UK biobank, and measured prediction using the area under the receiver operating curve (AUC). The standard model produced an AUC of 0.603 (SD=0.013), and the JASS weighted PRS produced and AUC of 0.621 (SD=0.013). Although the difference between both approaches was small (*P* = 0.095), it still suggests that neuroanatomical GWAS results could not only help identifying disorder associated variants but might also improve the predictive accuracy of mental disorder PRS.

### Supervised learning of disorder genetic risk based on MRI association

The strong enrichment of associations with ADHD and schizophrenia for variants significant in the multitrait MRI analysis confirmed some shared genetic factors between neuroanatomical traits and mental disorders. However, this enrichment was not informative regarding potential latent biomarkers involved in the genetics of these two outcomes. Indeed, the omnibus multitrait test is an eight degree of freedom test, whose significance is independent of the direction of the effects across the eight traits. That is, the multitrait test can capture equally a range of heterogeneous associations with the MRI phenotypes. In parallel, univariate MRI GWAS show neither concordant association with the disorder, nor enrichment for association (**Figs. 1-2**). Given these two results, we wondered whether a naïve supervised learning approach applied to the entire genome could recover potential linear combinations of the MRI phenotypes association statistics which could explain the enrichment for association with ADHD and schizophrenia. In practice, we performed a multiple regression where the Z-score of the disease was treated as the outcome and the Z-score of the eight MRI phenotypes at the same variants was use as predictor: *Z*_*disorder*_ = ∑_*i*=1…8_ *γ*_*i*_*Z*_*MRIi*_. We applied this approach using the top associated variant per locus and considered a range of threshold *P*_t_ on the *p*-value with the disorder (see **methods**). As showed in **Table S5**, we did not identify any significant linear combination able to predict the disorder genetic association after accounting for multiple testing. The adjusted *r*-square was very low for most of threshold considered (*r*-square < 0.004 for *P*_t_ = 5 x 10^-4^), although one subset showed nominal significance and slightly larger *r*-square for schizophrenia (*r*-square < 0.069, *P* = 0.028, for *P*_t_ = 5 x 10^-4^).

### Unsupervised learning to identify latent genetic pathways

The null result from the supervised learning suggested that the genetic relationship between mental disorder and MRI phenotypes could not be summarized at the genome-wide level and might instead be heterogenous across the genome. One possibility is that the neuroanatomical phenotypes depend on multiple genetic pathways with various effects on the disorders. Given the strong genetic correlation across MRI phenotypes (**Fig. 1**), we considered using an unsupervised learning approach, where variants were first grouped based on the clustering of the multitrait matrix of variants-trait association, and then tested jointly for association with the disorders. Intuitively, this approach assumes that there exist subsets of variants with similar multitrait association pattern (e.g., variants positively associated with two traits, and negatively associated with other traits), that encode distinct biological functions associated with mental disorders. **Figure 3** illustrates the clustering approach we implemented. In brief, we applied a spherical *k*-medoids clustering algorithm to the MRI association matrix, assuming a number of clusters between 2 to 12, resulting in a total of 77 possible clusters. For each cluster, we tested the association of a genetic risk score derived as *S* = ∑_*Ω*_ *W*_*i*_ *Z*_*i*_, where *W*_*i*_ and *Z*_*i*_ are a pre-specified weight and the Z-score of variant *i* in the disorder GWAS, and *Ω* is a set of variants within the cluster. The weight of each variant was defined so that variants close to the centroid of the cluster had large positive weight and those distant from the centroid had null or negative weights (**Figure 3** and **methods**).

**Figure 3.**
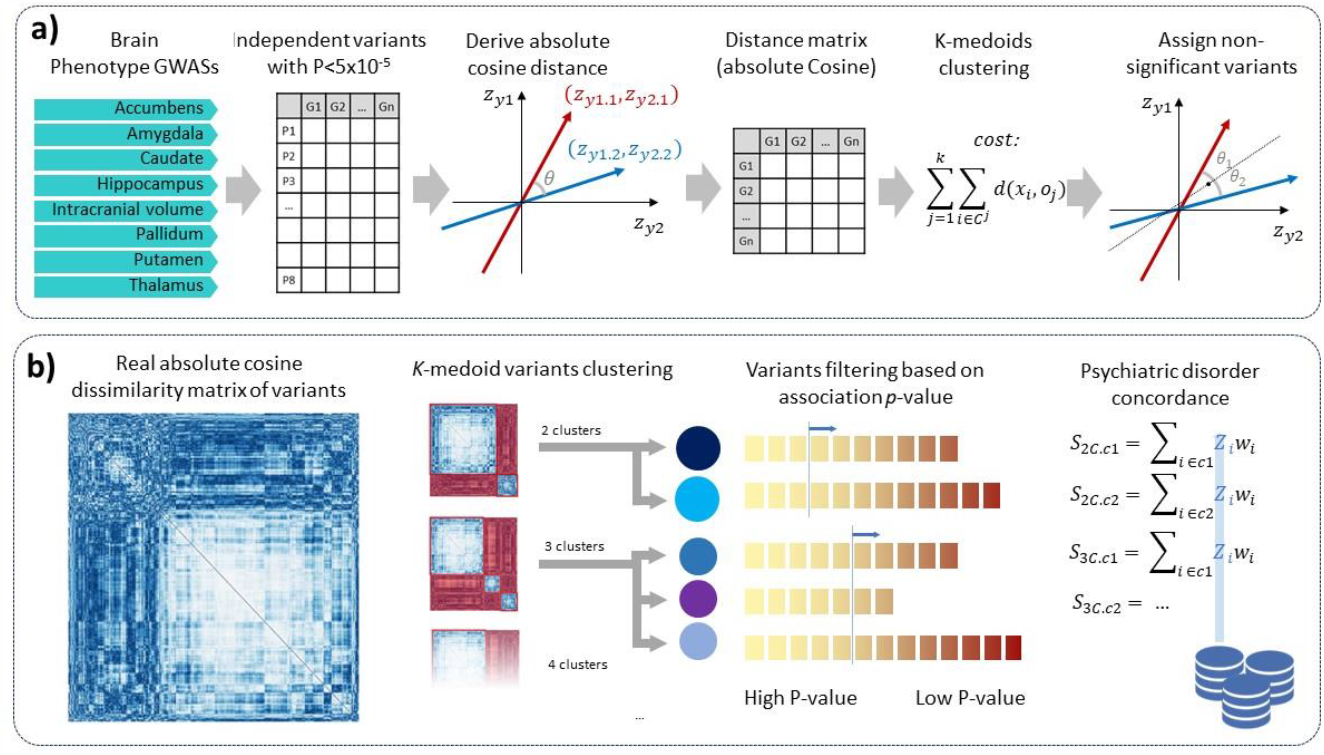
Clustering approach. We used univariate brain phenotypes association signal to build cluster of variants displaying similar multitrait association pattern. Panel **a)** illustrates the clustering pipeline. Top independent associated variants are selected based on their univariate and multivariate association (P < 5 x 10^-5^). A dissimilarity matrix is derived based on the absolute cosine distance. *K*-medoids clustering is applied on this distance matrix which minimize a cost function depending on the distance between each point *x* and a medoids *o*. Finally, independent non-significant variants (P > 5 x 10^-5^) are assigned to cluster *a posteriori*. Panel **b)** displays the real absolute cosine dissimilarity matrix derived from the top 479 variants. The *K*-medoids clustering was applied to this matrix assuming two and 12 clusters. Variants from each cluster are then selected based on their *p*-value for association with the brain phenotypes, and the resulting sets evaluated for enrichment in concordant effects with each of the six disorders using a weighted Z-score approach.

**Figure 4** presents the association between the top cluster and each of the six disorders, while filtering out variants based on their multitrait association *p*-value. We observed significant associations for the two disorders also identified in the JASS enrichment analyses. One cluster was significantly associated with ADHD (min *P* = 3.0 x 10^-9^) and another one with schizophrenia (min *P* = 2.0 x 10^-9^). There was no significant association with the other disorders. The top ADHD associated cluster was observed for a two-cluster model and included 939 independent variants (**Table S6**). The strongest association was observed when using a JASS *p*-value threshold of 1.0 x 10^-5^ resulting in a subset of 104 variants. The top schizophrenia associated cluster was observed when using an eight-clusters model and included 349 independent variants (**Table S7**). The strongest association was observed when using a JASS *p*-value threshold of 3.1 x 10^-5^ resulting in a subset of 60 variants. Note that there is substantial overlap of variants across the 77 clusters (e.g., cluster 1 from the two-clusters model necessarily overlap with some of the clusters from the three and more-clusters models, see **Fig. 3**). As showed in **Figures S3**, clusters overlapping with the best schizophrenia and ADHD clusters display enrichment approximately proportional to the overlap, confirming that these top clusters capture most of the association with the disorders. Finally, in line with the correlation analyses (**Fig. 1**), the application of the *S* test on subsets of variants selected based on univariate MRI association signal did not identify any association (**Figure 4**), except for Pallidum volume and ADHD (*P* = 8.19 x 10^-7^), that just reach the Bonferroni corrected significance threshold.

**Figure 4.**
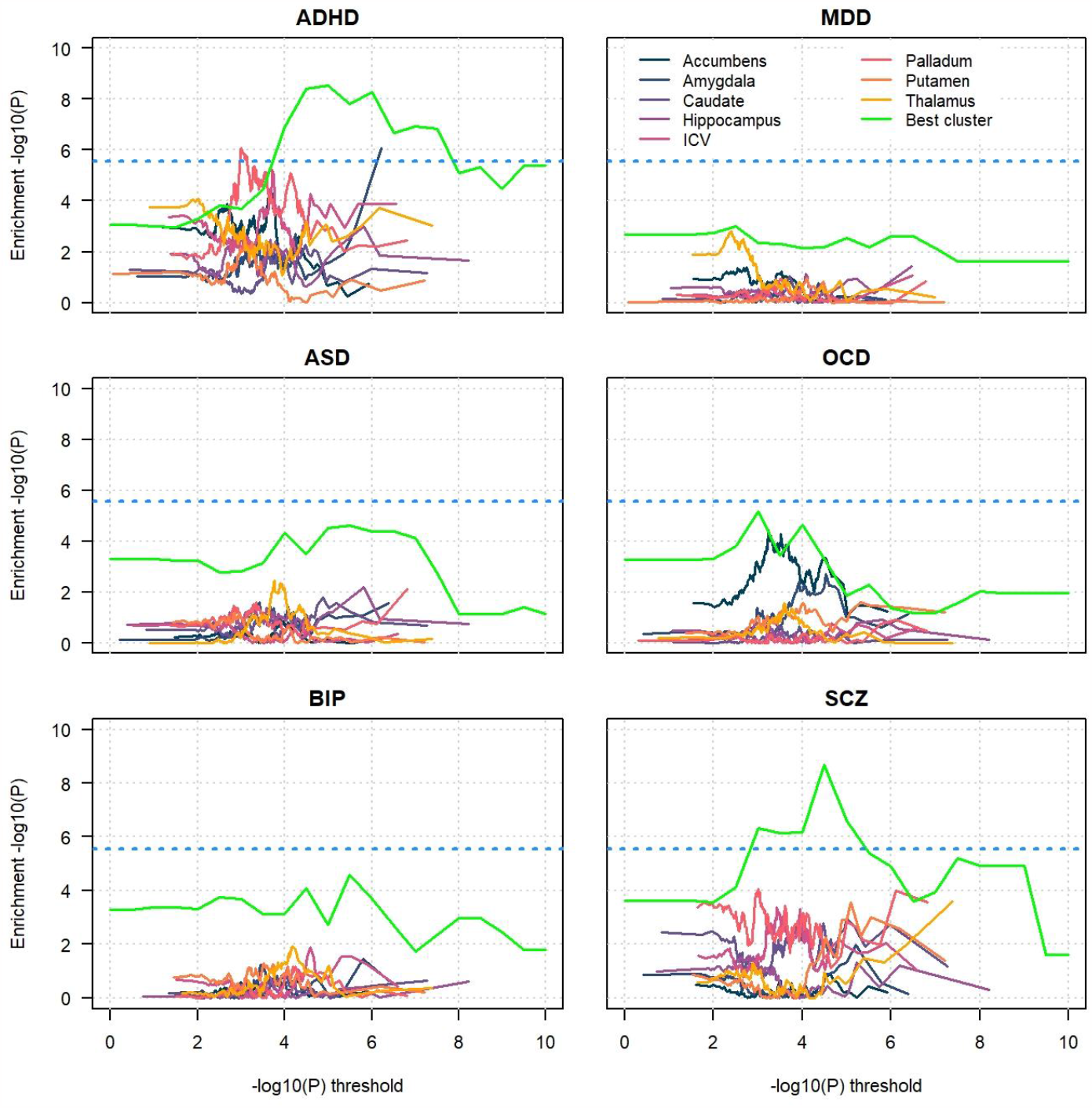
Concordance of effect between brain phenotypes and psychiatric disorders. Enrichment for concordant genetic associations signal between MRI phenotypes and each of the six psychiatric disorders: attention-deficit/hyperactivity disorder (ADHD), autism spectrum disorder (ASD), bipolarity (BIP), major depressive disorder (MDD), obsessive-compulsive disorder (OCD) and schizophrenia (SCZ). Concordant association for a set of variants corresponds to a case where, given the coded allele are defined as the increasing phenotype allele, the association with the disorder for those variants are consistently positive or consistently negative. For univariate MRI phenotypes, concordance was derived over subsets of independent variants selected based on their univariate association *p*-value. For clusters, we selected variants based on their multivariate MRI *p*-value. The X axis displays the *p*-value threshold. The Y axis represent the -log10(*p*-value) of the concordance obtained from a test of the weighted sum of disorder’s Z-score. The dash blue line shows the Bonferroni corrected significance threshold. The green line shows the enrichment for the most associated cluster.

### Characteristics of the latent neuroanatomical phenotypes

We mapped variants within each of the two clusters to their nearest genes and conducted a functional annotation overrepresentation analysis using DAVID^37^ (**Table S10**). The ADHD associated cluster was poorly specific, including a large number of variants all over the genome (**Fig. S3**). It showed highly significant overrepresentation of annotations but with low fold-enrichment (**Fig. S4, Table S8**). Largest fold-enrichment (5.28, Bonferroni corrected *P*=6.7 x 10^-5^) was observed for *protein phosphorylation*. The most significant one was for *Polar residues* (min *P* = 1.1 x 10^-10^), but enrichment was very modest (average = 1.7). The top significant annotation for the schizophrenia associated cluster was *neurotrophins signaling pathway* (Bonferroni corrected *P* = 2.2 x 10^-3^), which displays a very strong enrichment (up to 27.06, **Fig. S5, Table S9**), and was the best candidate mechanism underlying the shared genetics between MRI phenotypes and schizophrenia. Peack significance for that pathway strongly coincides with the association with schizophrenia. Moreover, recent meta-analysis suggests that neurotrophins might play a role in the pathophysiology of schizophrenia^38–40^ and it has been reported as a potential biomarker for brain function, structure and cognition^41^. The neurotrophins pathway overrepresentation was highly specific to this cluster, and we found no similar enrichment from top MRI variants after excluding those within that cluster. Overrepresented annotations from the two clusters, on top of those previously cited, included *kinase, EGFR tyrosine kinase inhibitor resistance*, and *Serine/threonine-protein kinase*. Several of these annotations have also been discussed in schizophrenia studies^42–44^, suggesting that other mechanisms might be at play. **Figure 5a** summarizes the annotation mapped to the two clusters along potential evidence for association with schizophrenia and ADHD extracted from a quantitative literature search.

**Figure 5.**
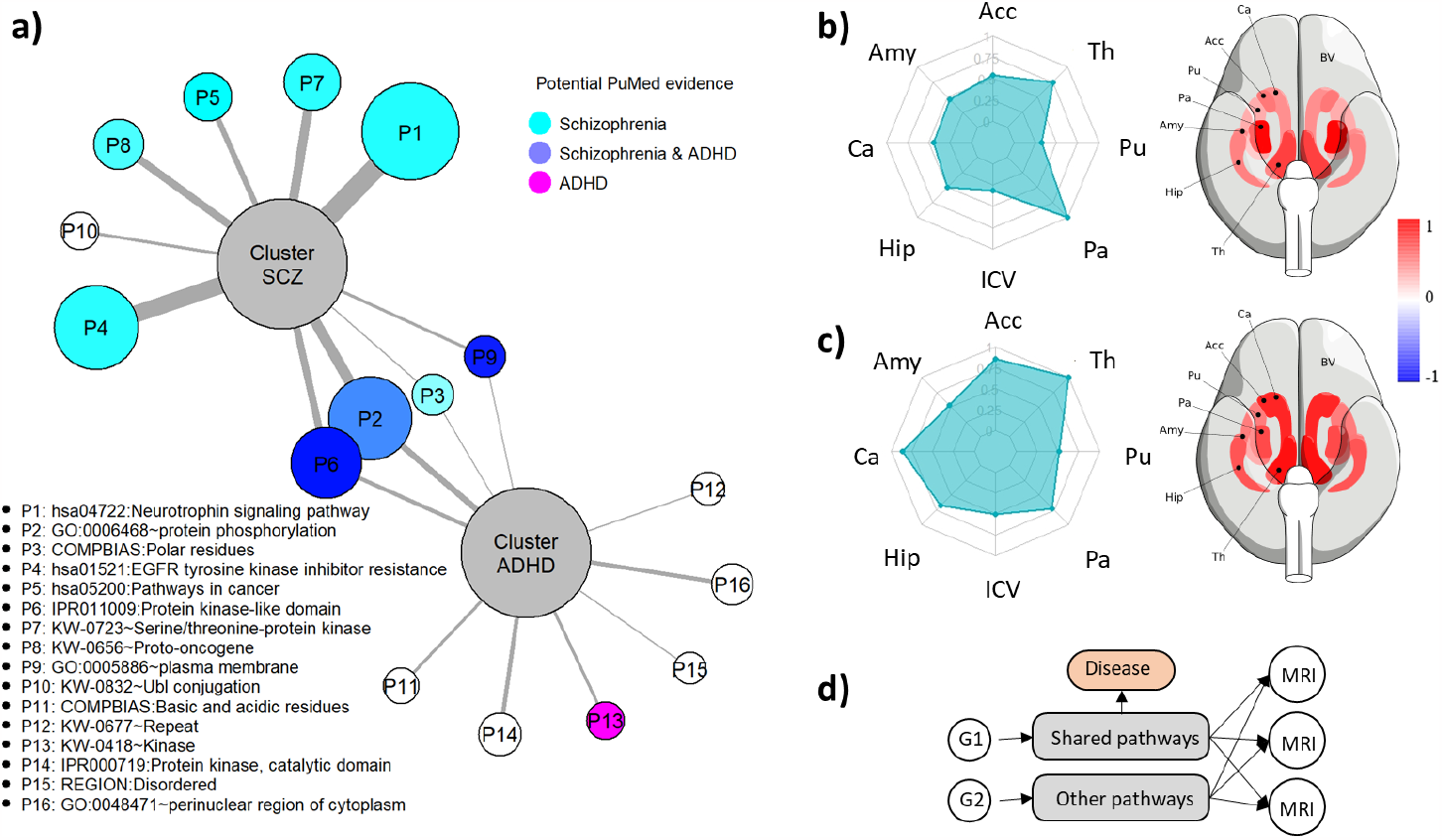
Overview of ADHD and schizophrenia associated clusters. **a)** Summary of the functional annotation analysis for the ADHD and schizophrenia clusters. Edge thickness and vertex size of the annotation are proportional to the fold enrichment. Colors of the vertex indicates potential evidence for a role in schizophrenia and ADHD from PMC search. Panels **b)** and **c)** present the weights of the cluster with the highest concordant signal for ADHD and schizophrenia (SCZ), respectively. Left panels present the absolute value of the weights with a radar plot. Right panels present the weights plotted on the corresponding brain area. These weights are the standardized Z-scores of the medoid for each MRI phenotypes: accumbens volume (Acc), amygdala volume (Amy), caudate volume (Ca), hippocampus volume (Hip), intracranial volume (ICV), pallidum volume (Pa), putamen volume (Pu) and thalamus volume (Th). **d)** Visual representation of the hypothetical mechanisms. The genetic component of MRI phenotypes involves multiple biological pathways, some of which being associated with an increased risk of mental disorders.

By construction, clusters inferred from the spherical *k*-medoids capture association effects in a specific direction in the multidimensional MRI space. The weights on each phenotype from each cluster can therefore be used to form a composite latent variable associated with the genetics variants (**Fig. 5b-c**). We computed this latent variable for the cluster associated with schizophrenia using the 20,744 UKB participants. A GWAS analysis of this variable identified 12 associated loci (**Figure S6** and **Table S11**), five of them not identified by the univariate MRI GWAS: rs9874537 (*P* = 4.5 x 10^-8^, min *P*_MRI_ = 3.4 x 10^-5^), rs768519054 (P = 4.6 x 10^-8^, min *P*_MRI_ = 3.4 x 10^-5^), rs298619 (*P* = 4.7 x 10^-10^, min *P*_MRI_ = 3.3 x 10^-7^), rs12762089 (*P* = 7.0 x 10^-9^, min *P*_MRI_ = 2.8 x 10^-4^), and rs28678082 (*P* = 3.4 x 10^-8^, min *P*_MRI_ = 4.8 x 10^-7^). Although several of these variants can be mapped to genes related to schizophrenia (e.g. AUTS2^45,46^ and ABCC5^47^), there was limited direct association with the disorder. Moreover, repeating the DAVID^37^ analysis on the top associated variants at various threshold did not point toward any significant annotation enrichment, including neurotrophins in particular. This is in agreement with the supervised clustering analysis that did no identified linear combination of phenotype relevant for the genetic of any of the disorder studied. It suggests that the MRI phenotypes studied were more likely influenced by specific genetic mechanisms shared with schizophrenia, two of them identified in our clustering analysis, rather than on a mediation path of the disorder (**Fig. 5d**).

## Discussion

Neuroimaging technologies hold great promise to link specific symptoms of mental health disorders to abnormal patterns of brain activity. Characterizing these links can help understanding the disorder etiology and ultimately improve diagnostic and suggest new classifications of patients to implement more personalized treatments. However, as other candidate biomarkers, neuroimaging data can suffer from multiple issues, including confounding^48^ and reverse causation^49^. In this study we showed that multivariate neuroimaging genomics approaches can provide a compelling framework to circumvent these limitations. Using a set of eight MRI phenotypes and six common mental disorders, we showed that a simple multitrait genetic analysis of MRI phenotypes can be much more informative on the genetics of disorder than any single univariate analysis. The proposed screening identified more MRI associated variants and identified enrichment for association with ADHD and Schizophrenia, when univariate MRI convey no information on the genetic risk of any of the six disorders considered. Our investigation of potential latent genetic variables underlying MRI phenotypes associated with the disorders did not find any compelling evidence when using a supervised genome-wide based approaches. Conversely, more advance modeling based on multitrait association clustering identified sets of variants underlying MRI phenotypes that were relevant for both outcomes. Our *in silico* functional annotation analysis further suggested the presence of a latent genetically-driven variable involved in the neurotrophins pathway influencing both the brain volume studied and schizophrenia^38,39,41^. This result opens paths for the inference of latent MRI variables, and ultimately for biomarker development and mediation-effects analyses.

This study has also some limitations. First, we used a set of eight MRI phenotypes without prior evaluation of their relevance towards the disorders considered. Future work using an optimal set of univariate phenotypes for a given disorder, or simply a much larger set of brain phenotypes, might maximize the detection of new association and help inferring more refined latent variables. The proposed framework might also be used along other approaches, including for example machine learning to extract MRI features of interest^50^. Second, we used an omnibus test to conduct our multitrait test. We previously showed that this fairly simple and standard approach performs well in multitrait GWAS analyses^51^. However, many alternatives have been proposed for the analysis of brain imaging data^8^. Comparing multivariate methods is out of the scope of this work, but we appreciate that other methods might further boost detection in new associations. Third, there exists hundreds of methods to perform data clustering. Here we used a spherical *k*-means as we believe its properties fitted well with the objective of identifying variants displaying similar multitrait association direction. The identification of two clusters of interest confirms the relevance of this approach. However, future work might consider alternative clustering methods. Fourth, we used mental disorder GWASs that typically use a broad disorder definition to maximize sample size. It is well accepted that mental disorders such as schizophrenia^52^ are hard to diagnose and even if the disorder is diagnosed, it can be divided into subtypes. Studying those subtypes might help refining potential links with MRI phenotypes. Large-scale GWAS of such disorder subtypes remain very sparse, but the proposed approach can easily be applied to those data as they become available.

Overall, the results of this study suggest that studying MRI phenotypes using a multivariate analysis approach can enhance the understanding of the links between these phenotypes and psychiatric disorders. The use of genetic variants identified through multivariate analysis and latent genetic variables derived from these variants can improve the detection of concordant signals for psychiatric disorders and opens paths for the development of biomarkers for the disorders.

## Acknowledgment

This work was supported by the Fondation pour la Recherche Médicale (ECO202106013759). It has been conducted as part of the INCEPTION program (Investissement d’Avenir grant ANR-16-CONV-0005). This research has been conducted using the UK Biobank Resource under Application Number 18584.

## Method

## Neuroanatomical brain phenotypes and mental disorder GWAS

We conducted genome-wide association studies (GWAS) of eight regional brain volumes: intra-cranial, accumbens, amygdala, caudate, hippocampus, pallidum, putamen and thalamus, on a subset of 20,744 unrelated UK Biobank participants (10,861 females, 9,883 males). Brain phenotypes were derived from raw MRI (magnetic resonance imaging) data using the *Freesurfer* software^53^. The phenotypes were selected based on a recent study from our group^26^. The GWAS included 11,993,198 variants either genotyped or imputed and was conducted using Plink2^27^ and standard quality control parametrization. We filtered out variants with imputation info score smaller than 0.8, missing rate above 0.05, minor allele frequency (MAF) below 0.1%, or Hardy-Weinberg equilibrium test below 1.0 x 10^-6^. We also removed individuals with missing rate larger than 0.1, or kinship relatedness larger than 0.025. Genomic control value (*λ*_*GC*_) ranged from 1.058 (putamen volume GWAS) to 1.070 (thalamus volume GWAS) (**Fig. S1**). For comparison purposes, we also used publicly available GWAS results for the same phenotypes from the ENIGMA consortium^30^. ENIGMA discovery sample included 13,171 individuals and approximately 7.5 million variants, with 6 million variants overlapping with the UK Biobank GWAS. Genetic association with brain phenotype were compared with GWASs of six mental disorders: attention-deficit/hyperactivity disorder (ADHD), autism spectrum disorder (ASD), bipolar disorder (BIP), major depressive disorder (MDD), obsessive-compulsive disorder (OCD) and schizophrenia (SCZ). GWAS from these disorders were conducted by the Psychiatric Genomic Consortium (PGC)^54–59^ and did not include UK Biobank participants (**Table S4**).

## Genetic correlation

Genome-wide correlations between brain phenotypes and psychiatric disorders were estimated using the LD-score python package^33,34^. Correlations based on top associated variants were done following the approach used by Pickrell et al^60^. Briefly, for each GWAS *j*, we selected the top associated variants per independent locus (see the next section for the definition of independent loci), extracted the association coefficient at those variants for the other GWAS *l* ≠ *j*, and derived the correlation between coefficient *ρ*_*jl*_ = *cor*(*β*_*j*_, *β*_*l*≠*j*_) with *β*_*j*_ the vector of genetic variants effect on phenotype j. Note that this result in an asymmetric matrix with *ρ*_*jl*_ potentially different from *ρ*_*lj*_, as the variants selected for the index GWAS *j* might differ from the variants selected for GWAS *l*. In the case of phenotype to phenotype correlation using independent variants, we accounted for bias introduced by phenotypic correlation and sample overlap using the following formula^51^:

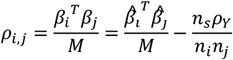

With *ρ*_*i,j*_ the corrected phenotypic covariance, *β*_*i*_ and *β*_*j*_ the vector of genetic effects for the pair of phenotypes i and 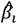 and 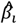 their respective estimates, *M* the number of variants, *n*_*i*_ and *n*_*j*_ the respective sample size for phenotypes *i* and *j, n*_*s*_ the sample overlap and *ρ*_*Y*_ the genome-wide phenotypic covariance.

## LD-region based comparison

Except if specified otherwise, comparisons across GWAS results were done using the top associated variants from independent loci defined based on linkage disequilibrium (LD) blocks^61^. Briefly, the genome was split into 1,703 loci using 1000 Genomes Phase 1 dataset as a reference panel. Overall, 1668 loci had GWAS summary statistics for at least one phenotype. Focusing on the 7,099,802 variants with data on both MRI and disorders, loci included 4256 variants on average, with a maximum of 9,859 and a minimum of 16. Using those independent loci for comparison purpose avoid overcounting association signals for correlated variants and addresses the issue of missing summary statistics (e.g., when the top associated variant for one phenotype is not available for another phenotype).

## Multitrait GWAS analysis

Multitrait analyses were performed using JASS^29^, a polyvalent Python package that allows for the computation of various joint test from GWAS summary statistics. Here, we applied an omnibus test, where a joint statistic *T*_*omni*_ is derived as: *T*_*omni*_ = *z*^*T*^*Σ*^−1^*z*, where *z* = (*z*_1_, *z*_2_, …, *z*_*k*_) is a vector of *k* Z-scores (for *k* GWAS results), derived from the summary statistics as 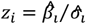 where 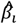 and 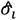 are the estimated regression coefficient and its standard error for study *i* ; and *Σ* is the covariance matrix between the Z-scores under the null. The latter covariance, *Σ*, is approximated by the intercept covariance matrix from multitrait LDscore regression^33,34^ applied to the GWAS summary statistics. Under the null hypothesis of no association between the variant tested and any of the *k* phenotypes tested jointly, *T*_*omni*_ follows a *χ*^*2*^ distribution with *k* degrees of freedom. As for univariate GWAS analyses, results comparisons were conducted using the top associated variants from predefined independent loci.

## Enrichment for association with psychiatric disorders

We estimated whether genetic variants selected based on univariate and multivariate association results with the MRI phenotypes were enriched for association signal with the psychiatric disorders. Enrichment was conducted for each brain phenotype-psychiatric disorder pair using the top associated variants per locus, starting with 1,668 top variants with the lowest *p*-values, then 1660 and iteratively removing 10 variants, going down to the top 10 independent variants with the lowest *p*-value. We tested the enrichment using a joint association statistic between selected variants and each disorder and derived as: *S*_*u*_ = ∑_*i*=1…*n*_ *χ*^2^_*i*_, where *χ*^2^_*i*_ = (*χ*^2^_1_, …, *χ*^2^_*n*_) is a vector of *n* chi-squared (for *n* variants). Under the null hypothesis of no association, *S*_*u*_ is expected to follow a chi-squared distribution with *n* degrees of freedom. However, this test itself is not necessarily a good indicator of potential enrichment. Indeed, because of the large sample size of the disorder GWAS and polygenicity, the association Z-scores do not follow a standardized normal distribution, but instead display a variance larger than 1. As a result, random sets of variants display more significant association with the outcome than expected by chance from null data. Modeling the null distribution of *S*_*u*_ from a simple polygenic model based on the estimated variance of the Z-score was unsuccessful (data not shown). Instead, we derived *S*_*u*.*null*_, an empirical distribution of *S*_*u*_ under the null based on the random sampling of *N*_*r*_ variants. This distribution was then used to derive the *p*-value for enrichment as: 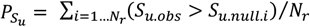. Due to computational cost, we limited the number of samplings, *N*_*r*_ to 10,000,000, so that the minimum *p*-value that can be derived equals 1 x10^-7^.

## Polygenic risk score

We investigated the potential of using association results from the multitrait brain phenotypes analysis to improve the predictive power of schizophrenia polygenic risk models. As multitrait analysis does not produce a signed statistic, it cannot be used to directly produce PRS weights. Instead, we relied on a the SbayesRC^36^ approach, a polyvalent method that allow to select the variants for polygenic risk score analyses using binary and continuous annotations. We applied SbayesRC to select variants from the Schizophrenia GWAS with and without the multitrait signal annotation. Those variants were passed to PRSice^62^ to compute the PRS for both models. For the annotated PRS, we generated a continuous annotation from the eight degree of freedom chi-square statistic from the multi-trait analysis. To limit the impact of extreme statistics, we set the maximum value of the annotation to the value of the 99.9% highest percentile (*χ*^2^ = 40).

The two sets of weights for *M* variants, *W* = (*W*_1_ … *W*_*M*_), were used to compute a polygenic score defined as *PRS* = ∑_*M*_ *W*_*i*_*G*_*i*_ in a 585 Schizophrenia cases and 9,396 controls from the UK Biobank. The performance of each PRS was evaluated using the area under the receiver operating curve (AUC), and compared using a one-sided Z-score test: 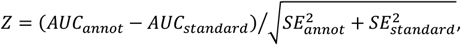 where *AUC*_*annot*_ and *AUC*_*standard*_ are the AUC from the standard and annotated PRS, and 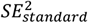 and 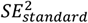 are their respective variance. Under the null, *Z* is expected to follow a standard normal distribution.

## Supervised learning of disorder genetic risk based on MRI association

We performed a supervised learning of the disorder GWAS summary statistics using MRI GWAS as predictors. The goal was to determine whether association of variants with the disorder can be predicted based on a linear combination of the MRI association statistics. In practice, we used a multiple regression model as : *Z*_*disorder*_ = ∑_*i*_=1…8 *γ*_*i*_*Z*_*MRIi*_ where the outcome *Z*_*disorder*_ is the vector of disorder variants Z-score and *Z*_*MRIi*_ is the vector Z-score for MRI phenotype*i* for those same variants. The regression was applied for ADHD and schizophrenia using the top associated variant per locus from each disorder, including variants significant over a *p*-value threshold we varied from 1 to 2.5 x 10^-10^. For each variant set, we derived the R-squared and the model *p*-value.

## Assessing concordant associations between psychiatric disorders and brain phenotypes

We assessed the concordance of effect’s directions between brain phenotypes and each disorder, that is, whether the alleles associated with an increase in phenotypic value were consistently associated with either a decrease or an increased risk of the disorders. As for the enrichment analysis, we used independent genetic variants selected based on their *p*-value for association with the brain phenotypes, using the most associated variant per LD-locus. Concordance was tested sequentially starting with the top 1,668 associated variants from each of the 1,668 LD-regions and decreasing that initial set, then the 1660 most associated variants, then decreasing by incrementally removing the 10 less significant variants, thus testing a total of 170 sets. There are various approaches to test for concordance of effect. Here we used a weighted sum of Z-scores statistics defined as *S*_*s*_ = ∑_*q*=1…*Q*_ *Z*_*q*_ × *W*_*q*_ with *Z*_*q*_ the Z-score of variant *q* for the disorder and *W*_*q*_ the Z-score for the phenotype tested. Under the null, the variance of *S*_*s*_ is *var*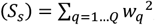, and a test for concordance of effect can be derived as 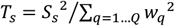 . Under the null, *T*_*s*_ follow a one degree-of-freedom chi-square distribution.

## Enrichment for concordant associations after clustering of variants based on multitrait association

We conducted a clustering analysis to examine whether subsets of variants selected based on multi-trait association similarity display concordant association with disorders. The clustering was performed on variants with an association *p*-value under 5x10^-5^ for either the multitrait test or the univariate brain phenotype test using a spherical *k*-medoids method, where the clustering is applied to the absolute Cosine distance between data points. The cosine distance is defined as 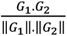 with *G*_*i*_ a vector containing the coordinates of SNP *i*. These coordinates are the first five principal components computed from a principal component analysis we conducted on the suggestive variants we selected. Briefly, the absolute cosine distance between two points measures whether the two associated multidimensional vectors are pointing in the same direction.

The *k*-medoids clustering was conducted using the R Cluster^63^ package. The *k*-medoids requires to pre-specify the number of clusters. As the number of relevant clusters is unknown, we used a systematic screening approach, applying the clustering while assuming 2 to 12 clusters, resulting in a total of 77 (overlapping) clusters. All variants not used in the clustering (i.e., with a *p*-value above 5 x10^-5^) were assigned *a posteriori* to the closest cluster using again the absolute Cosine distance to the closest medoids (a specific variant output by the *k*-medoid analysis representing the center of each cluster along). However, to further increase the homogeneity of multitrait effect across variants, within clusters, we filtered out variants displaying an absolute cosine distance to the medoid above 0.5.

For each cluster, we computed the enrichment for association with each disorder with prior on the direction of effects using the same approach as for the concordant enrichment of brain phenotypes with a psychiatric disorder, that is: *S*_*c*_ = ∑_*i*_∈_*n*_ *Z*_*i*_*W*_*i*_ where *Z*_*i*_ is the Z-score for a disorder for variant *i*. The weight *W*_*i*_ was defined as followed: 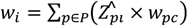 with 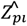 the Z-score of variant *i* for phenotype *p, P* the number of phenotypes and *W*_*pc*_ the Z-score of the medoid of cluster *c* for phenotype *p*. Under the null hypothesis of no enrichment, 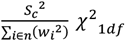 . For a threshold from 0 to 10 with increments of 0.5, we tested each cluster using SNPs with a -log10(minimum *p*-value) above the threshold and we kept the most associated cluster for each threshold. To avoid clusters with a signal led by a few SNPs, we applied two additional conditions. First, we only considered clusters with at least five SNPs above the threshold. Second, when a cluster is significant, we remove the three variants with the highest |Z-score| within the cluster and compute a second *S*_*signed*_ value. If it is not significant, it is removed from the analysis.

## Functional annotation overrepresentation analysis

We conducted a functional annotation overrepresentation analysis using DAVID^37^. First, each independent variant considered was mapped to its closest gene using the *closest* function in BEDtools^64^, and the gencode release 44 annotation database^65^ (**Table S6-7**). Genes were submitted to DAVID as a gene list and fold enrichment was derived over the complete DAVID Knowledge-base. Overrepresentation analysis was applied to subset of variants from the ADHD and schizophrenia associated cluster which multitrait association *p*-value threshold. For each cluster, we report the top 10 annotation pathways showing overrepresentation significant after Bonferroni correction (**Table S8-9**). To assess the relevance of the enrichment we observed with the neurotrophins signaling pathway results in the schizophrenia cluster, we ran two other analyses using top independent variants of the cluster analysis without the variants of the cluster of interest, i) without threshold on the multivariate *p*-value and ii) with a multivariate *p*-value threshold of 5 x 10^-5^. Finally, we conducted a quantitative PMC (PubMed Central) search for publications for each annotation term and the corresponding disorder. PMC search was conducted using the disorder in the [Abstract/Title], and the annotation term in the [Body-Key Term] (**Table S10** and **Figure 5a**).

## Latent brain phenotype inference from multitrait clustering for the GWAS and MR study

We conducted genome-wide association studies (GWAS) of the latent variables on the 20,744 UK Biobank participants used in the univariate MRI GWAS. For each participant *j*, a latent variable was computed as the cosine distance similarity between a vector *Y*_*j*_ = {*Y* _*j*,1_, …, *Y* _*j,p*_} and *W*_*c*_ = {*W*_1,*c*_, …, *W*_*pc*_}with *Y* _*j,p*_ the standardized phenotype *p* of participant *j* and *W*_*pc*_the Z-score of the medoid of cluster *c* for phenotype *p*. This medoid is computed as the medoid of the subgroup of variants with the highest association, i.e. the top cluster for schizophrenia using only variants with a -log10(multivariate p-value) over 4.5 (**Figure 4**). Values of *W*_*pc*_ are displayed in **Figure 5b-c**.

## Online resources

JASS : https://gitlab.pasteur.fr/statistical-genetics/jass_suite_pipeline

