## Supplementary figures for "Linking the genetic structure of neuroanatomical phenotypes with psychiatric disorders"

**Supplementary Material and Methods**

### Supplementary Figures

#### Figure S1. Univariate brain phenotype GWAS

Manhattan plot (left) and QQplot (right) for the genome-wide association studies of the eight brain phenotypes: accumbens, amygdala, caudate, hippocampus, intracranial volume, pallidum, putamen and thalamus. Lambda values equal 1.033, 1.042, 1.041, 1.039, 1.042, 1.036, 1.036, 1.043, respectively. The red dash line on the Manhattan plot indicates the standard genome-wide significance threshold of 5 x10^-8^.

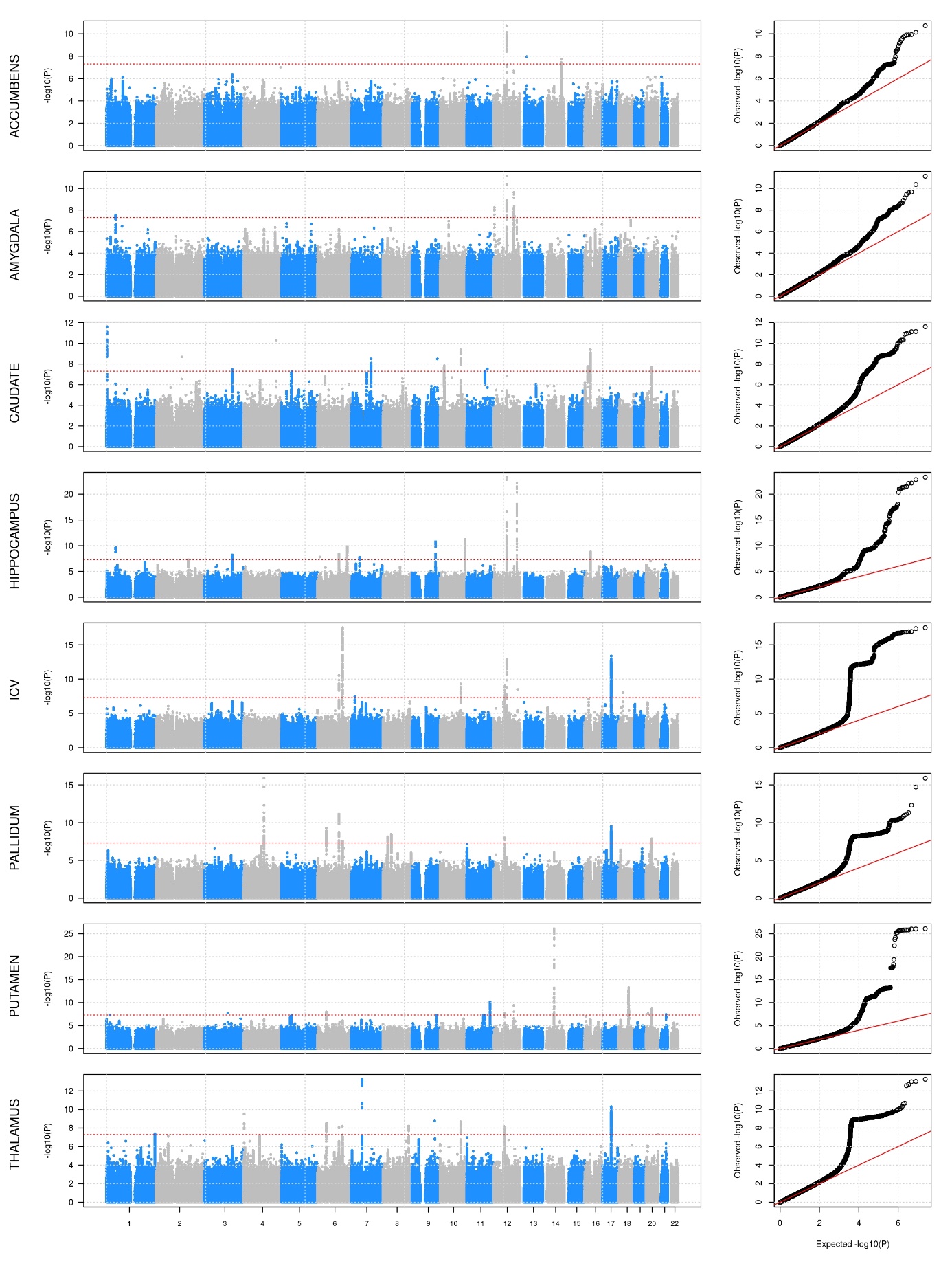

#### Figure S2. Multitrait GWAS analysis

Manhattan plot for the genome-wide association of the joint analysis of the eight brain phenotypes (accumbens, amygdala, caudate, hippocampus, intracranial volume, pallidum, putamen and thalamus) (a) and best univariate association across the eight univariate GWAS (b). QQplot for the joint test is presented in panel (c). The red dash line on the Manhattan plots indicates the standard genome-wide significance threshold of 5 x 10^-8^ applied for the joint test, and 6.25 x 10^-9^ applied for the best univariate GWAS. Panel (d) displays the quadrant plot of the top association per LD independent region (Ntotal = 1,668).

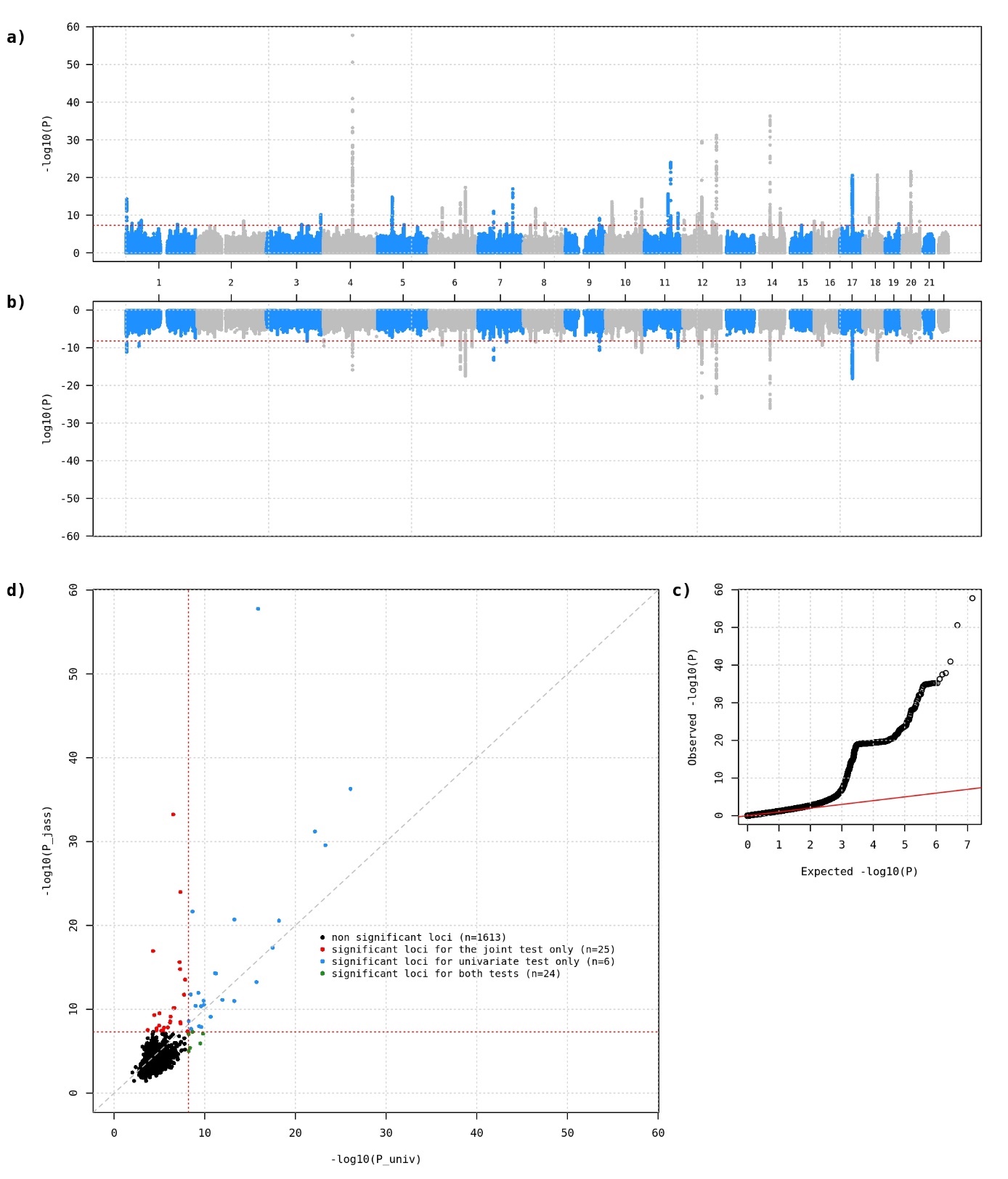

#### Figure S3. Enrichment for disease association across overlapping clusters.

Enrichment for concordant association with ADHD (left) and schizophrenia (right) for variants within the 77 (overlapping) clusters. Concordant association for a set of variants corresponds to a case where, given the coded allele are defined as the increasing phenotype allele, the association with the disorder for those variants are consistently positive or consistently negative. For each cluster, enrichment was derived over subsets of independent variants selected based on their multitrait MRI association *p*-value (X axis). The Y axis represent the -log10(*P*-value) of the enrichment. The cluster displaying the strongest enrichment with ADHD and schizophrenia, represented in Figure 4, are indicated by a dash line. The gradient of green for the other clusters indicates the proportion of variants from each cluster overlapping with that top cluster. The dash blue line shows the Bonferroni corrected significance threshold.

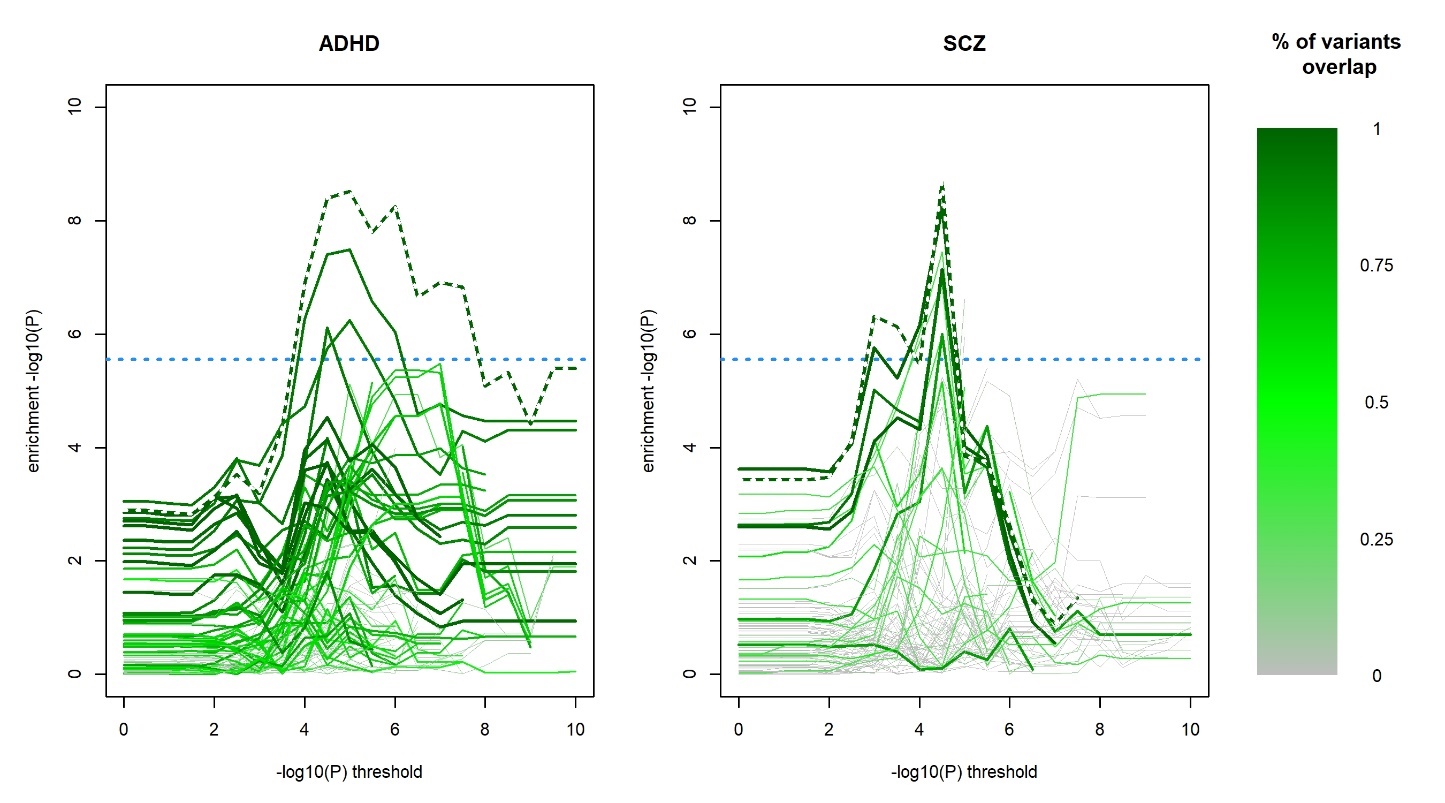

#### Figure S4. Functional annotation enrichment for the ADHD associated cluster

Functional annotation enrichment conducted using the DAVID web server on the nearest genes from each variant within the MRI cluster associated with schizophrenia. Enrichment was applied for all 939 genes from the 939 variants from the cluster, and on subset of genes selected based on their *p*-value for association from the multitrait MRI analysis (X axis). The figure presents the fold enrichment (Y axis) from the top annotations significant after Bonferroni correction. The size of each point is inversely proportional to the Bonferroni corrected *p*-value from DAVID. The green dashed line indicates the association of the cluster with schizophrenia.

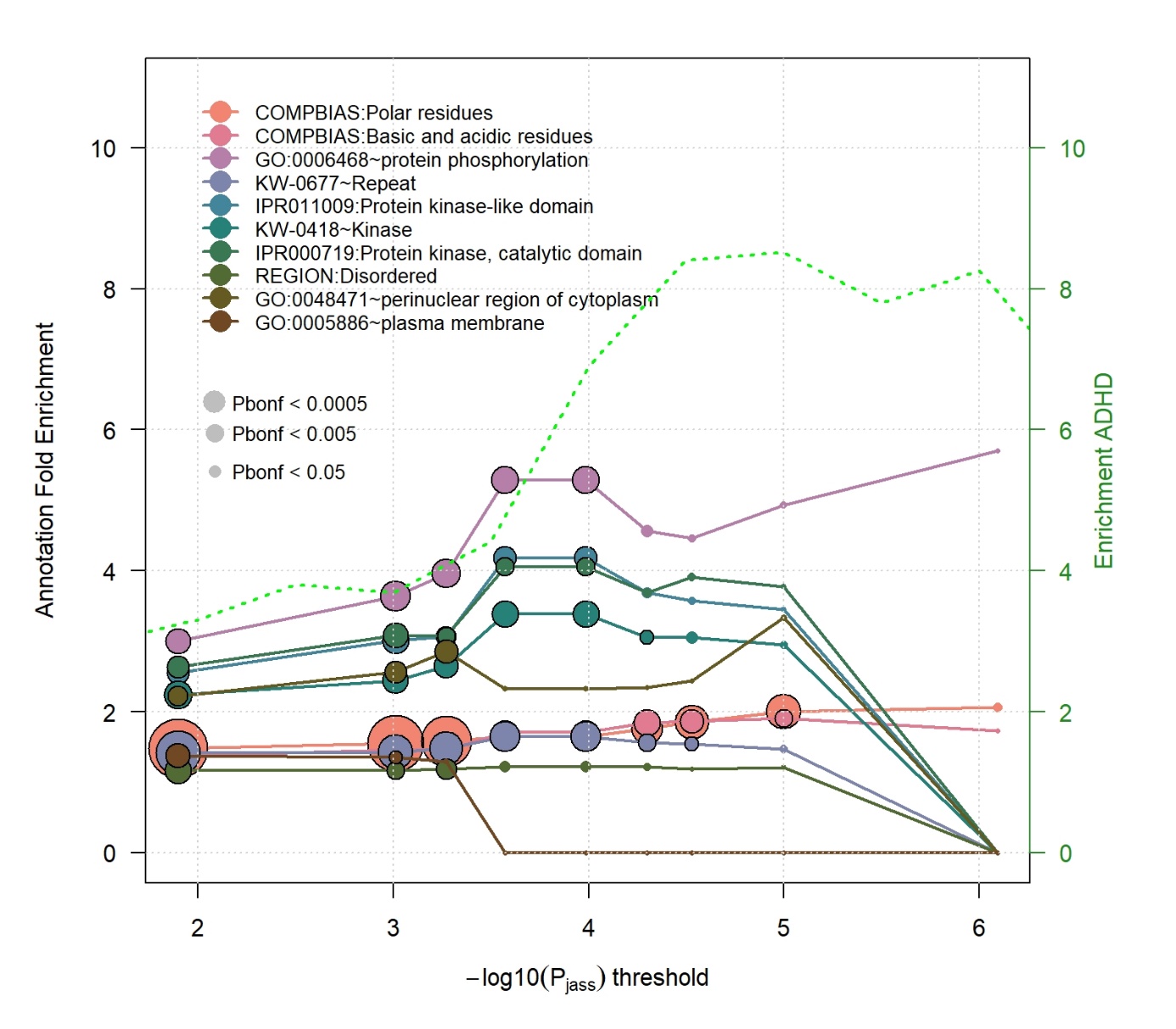

#### Figure S5. Functional annotation enrichment for the schizophrenia associated cluster

Functional annotation enrichment conducted using the DAVID web server on the nearest genes from each variant within the MRI cluster associated with schizophrenia. Enrichment was applied for all 349 genes from the 349 variants from the cluster, and on subset of genes selected based on their *p*-value for association from the multitrait MRI analysis (X axis). The figure presents the fold enrichment (Y axis) from the top annotations significant after Bonferroni correction. The size of each point is inversely proportional to the Bonferroni corrected *p*-value from DAVID. The green dashed line indicates the association of the cluster with schizophrenia.

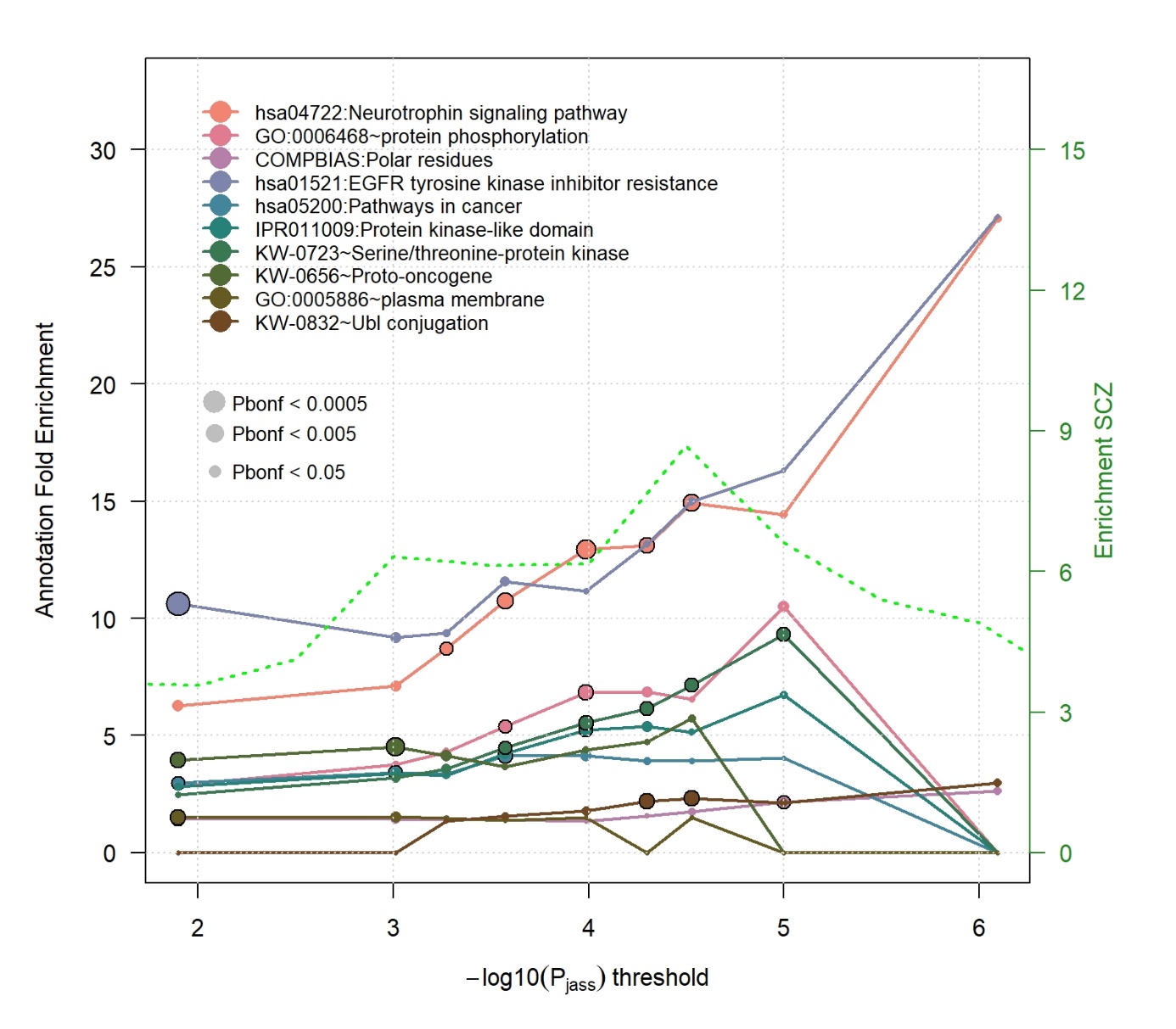

#### Figure S6. Latent variable GWAS for Schizophrenia

Manhattan plot (left) and QQplot (right) for the genome-wide association study of the latent variable derived from the cluster displaying the strongest association with schizophrenia. The red dash line on the Manhattan plot indicates the standard genome-wide significance threshold of 5 x 10^-8^. Lambda value derived from the QQplot equals 1.056.

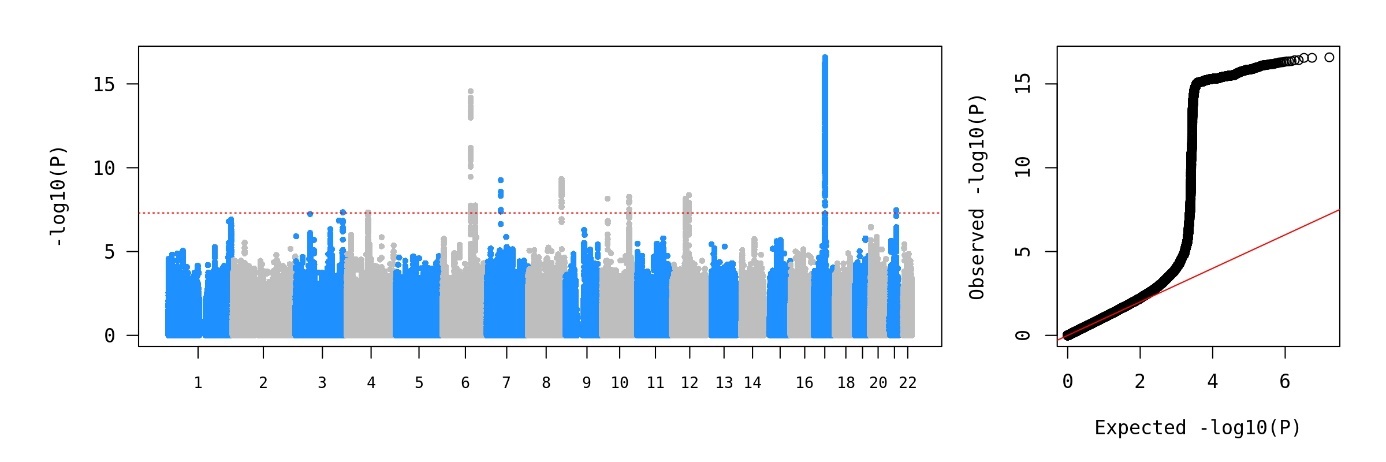
